# Exosomes from COVID-19 patients carry tenascin-C and fibrinogen-β in triggering inflammatory signals in distant organ cells

**DOI:** 10.1101/2021.02.08.430369

**Authors:** Subhayan Sur, Mousumi Khatun, Robert Steele, T. Scott Isbell, Ranjit Ray, Ratna B. Ray

## Abstract

SARS-CoV-2 infection causes cytokine storm and overshoot immunity in humans; however, it remains to be determined whether genetic material of SARS-CoV-2 and/or virus induced soluble mediators from lung epithelial cells as natural host are carried out by macrophages or other vehicles at distant organs causing tissue damage. We speculated that exosomes as extracellular vesicles are secreted from SARS-CoV-2 infected cells may transport messages to other cells of distant organs leading to pathogenic consequences. For this, we took an unbiased proteomic approach for analyses of exosomes isolated from plasma of healthy volunteers and SARS-CoV-2 infected patients. Our results revealed that tenascin-C (TNC) and fibrinogen-β (FGB) are highly abundant in exosomes from SARS-CoV-2 infected patient’s plasma as compared to that of healthy normal controls. Since TNC and FGB stimulate pro-inflammatory cytokines via NF-κB pathway, we examined the status of TNF-α, IL-6 and CCL5 expression upon exposure of hepatocytes to exosomes from COVID-19 patients and observed significant increase when compared with that from healthy subjects. Together, our results demonstrated that soluble mediators, like TNC and FGB, are transported through plasma exosomes in SARS-CoV-2 infected patients and trigger pro-inflammatory cytokine expression in cells of distant organs in COVID-19 patients.

**Importance:** Exosomes play an important role in intercellular communication by inducing physiological changes in recipient cells by transferring bioactive proteins. Little is known about exosomes from SARS-CoV-2 infected cells and their role in pathogenesis. Here, we have carefully examined and analyzed this aspect of SARS-CoV-2 infection. Our results uncovered the potential mechanisms by which SARS-CoV-2 communicates with other cells of distant organs and promotes pathogenesis. We expect to detect whether other factors are modulated in the presence of COVID-19 exosomes. Our exosomes related proteomic experiments prioritize after initial verification to further examine their role in SARS-CoV-2 associated other pathogenic mechanisms to target for therapeutic modalities.

## Introduction

Emergence of a novel severe acute respiratory syndrome coronavirus 2 (SARS-CoV-2) has precipitated the current global health crisis with many deaths worldwide. The World Health Organization announced COVID-19 to be a Public Health Emergency of International Concern and declared COVID-19 as a pandemic. SARS-CoV-2 is an enveloped virus containing a 29.9 kb positive-sense RNA genome. The virus genome contains at least ten open reading frames (ORFs). The first ORF (ORF1a/b), representing about two-thirds of the viral RNA, are translated into two large polyproteins, which are processed into 16 non-structural proteins (nsp1-nsp16), and some of them form the viral replicase transcriptase complex (1). The other ORFs of SARS-CoV-2 genome encode four main structural proteins: spike (S), envelope (E), nucleocapsid (N), membrane (M), and several accessory proteins of unknown functions.

The lung is a vital organ which supports blood oxygenation and decarboxylation necessary for aerobic life. Any insult including viral infection may impair this process and compromise survival. Lung immune responses and inflammatory processes are tightly regulated to maintain respiratory function. The lungs are highly susceptible in developing innate immune responses to viral infection, such as SARS-CoV-2. Although this viral envelop protein interacts with angiotensin-converting enzyme 2 (ACE2) present on many cell surfaces as a receptor, lung epithelial cells are probably the most susceptible cells for SARS-CoV-2 entry and replication causing human disease. Clinical observations indicate that severely ill COVID-19 patients develop extra-pulmonary tissue/organ dysfunctions, although viremia is not common. The presence of SARS-CoV-2 RNA in blood was reported in very low number of postmortem samples (2) and these observations are debatable (3). The pathophysiology of extra-pulmonary manifestations is not entirely clear but thought to occur in part to a dysregulated inflammatory response characterized by inhibition of interferon signaling by the virus, T cell lymphodepletion, and the production of pro-inflammatory cytokines, particularly IL-6 and TNFα (4, 5). Extra-pulmonary manifestations of SARS-CoV-2 associated disease have been documented in numerous organ systems including but not limited to cardiac, neurologic, hemostatic, kidney, and liver.

Exosomes (30–150 nm) are extracellular vesicles and play an important role in intercellular communication by inducing physiological changes in recipient cells by transferring bioactive lipids, nucleic acids, and proteins. Exosome vesicles are formed by the interior budding of endosomal membranes to form large multivesicular bodies (MVBs). Exosomes play an important role in cellular homeostasis and in the pathogenesis of major human diseases. Evidence suggested that exosomes carry materials from one cell to other cells for initiation and exaggeration of disease (6). Exosomes are also involved in viral spread, immune regulation, and antiviral response during infection (7–9). Little is known about exosomes of SARS-CoV-2 infected patients and their role in pathogenesis. In this study, we have observed that exosomes from plasma of COVID-19 patients harbor tenascin-C and fibrinogen-β to trigger inflammatory signal in distant cells.

## Results

### Isolation and characterization of exosomes from COVID-19 patient plasma

Patient information and samples used in this study are shown (Table 1). We have isolated exosomes from plasma of 20 COVID-19 patients and 8 healthy volunteers (normal). The size and purity of exosomes were examined by transmission electron microscopy (TEM) and observed spheres of heterogeneous size (30-70 nm particles) **(Figure 1, panel A)**. The presence of exosomal markers, CD63 and TSG101, was verified from the exosome preparations by Western blot analysis as described previously (10, 11) and the results from a representative blot is shown **(Figure 1, panel B)**. Exosome depleted serum (Invitrogen) was used a negative control and as expected, did not exhibit the presence or cross-reactivity for CD63 or TSG101 proteins. N1 and N2 of SARS-CoV-2 RNAs were undetected in exosome preparation from patients or healthy subject controls using a CDC recommended PCR kit.

**Table 1:** Sample information of COVID-19 patients

| **Parameters** |  | **Number** |
| --- | --- | --- |
| **COVID19 patients** |  | 20 |
| **Age** | >60 y | 11 |
|  | <60 y | 9 |
| **Sex** | Male | 8 |
|  | Female | 12 |
| **Deceased** | Yes | 7 |
|  | No | 13 |
| **D-dimer** | <0.5 µg/mL FEU | 1 |
|  | >0.5 µg/mL FEU | 15 |
|  | Unknown | 4 |
| **Location** | ICU | 20 |

**Figure 1:**
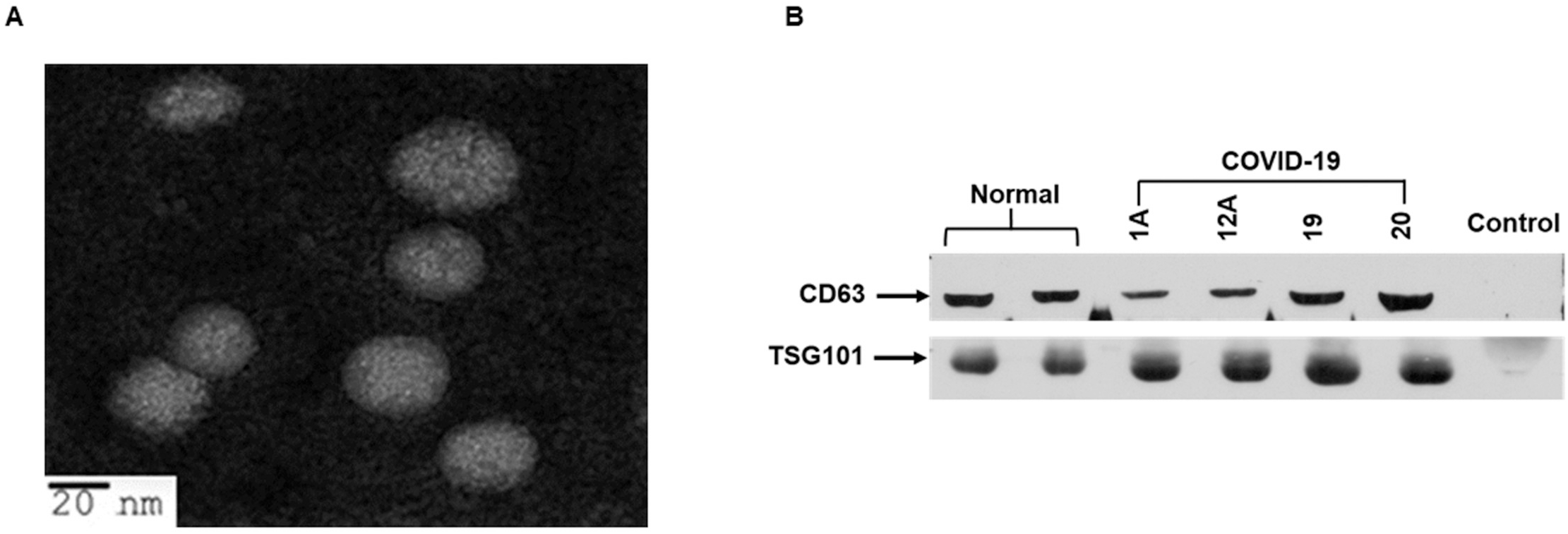
Scanning microscopy and characterization of exosomes from plasma of COVID-19 patients. **Panel A:** Representative transmission electron microscopy (TEM) image of exosomes isolated from COVID-19 patient plasma. The sample grids were screened under JEOL JEM-1400Plus Transmission Electron Microscope. The exosomes were round in shape with diameters of 30-70 nm. **Panel B:** Exosome lysates from plasma of normal and COVID-19 patients were subjected to Western blot analysis for detection of CD63 and TSG101 protein using specific antibodies. A representative image show results from normal and COVID-19 exosomes (1A, 12A, 19, 20). Exosome depleted serum was used as a negative control.

### Exosomes from plasma of COVID-19 patients harbor tenascin-C and fibrinogen-β

Little is known about the proteome profile of the exosomes from COVID-19 patients. Using unbiased proteomic approach, the mass spectrometry analysis identified 1,637 proteins. We shortlisted 163 proteins having more than five spectra counts and at least twofold changes as compared to plasma exosomes from healthy volunteers **(Figure 2, panels A and B)**. Tenascin-C (TNC) and fibrinogen-β (FGB) were identified as the two significantly enriched molecules in exosomes from COVID-19 patients relative to normal exosomes.

**Figure 2:**
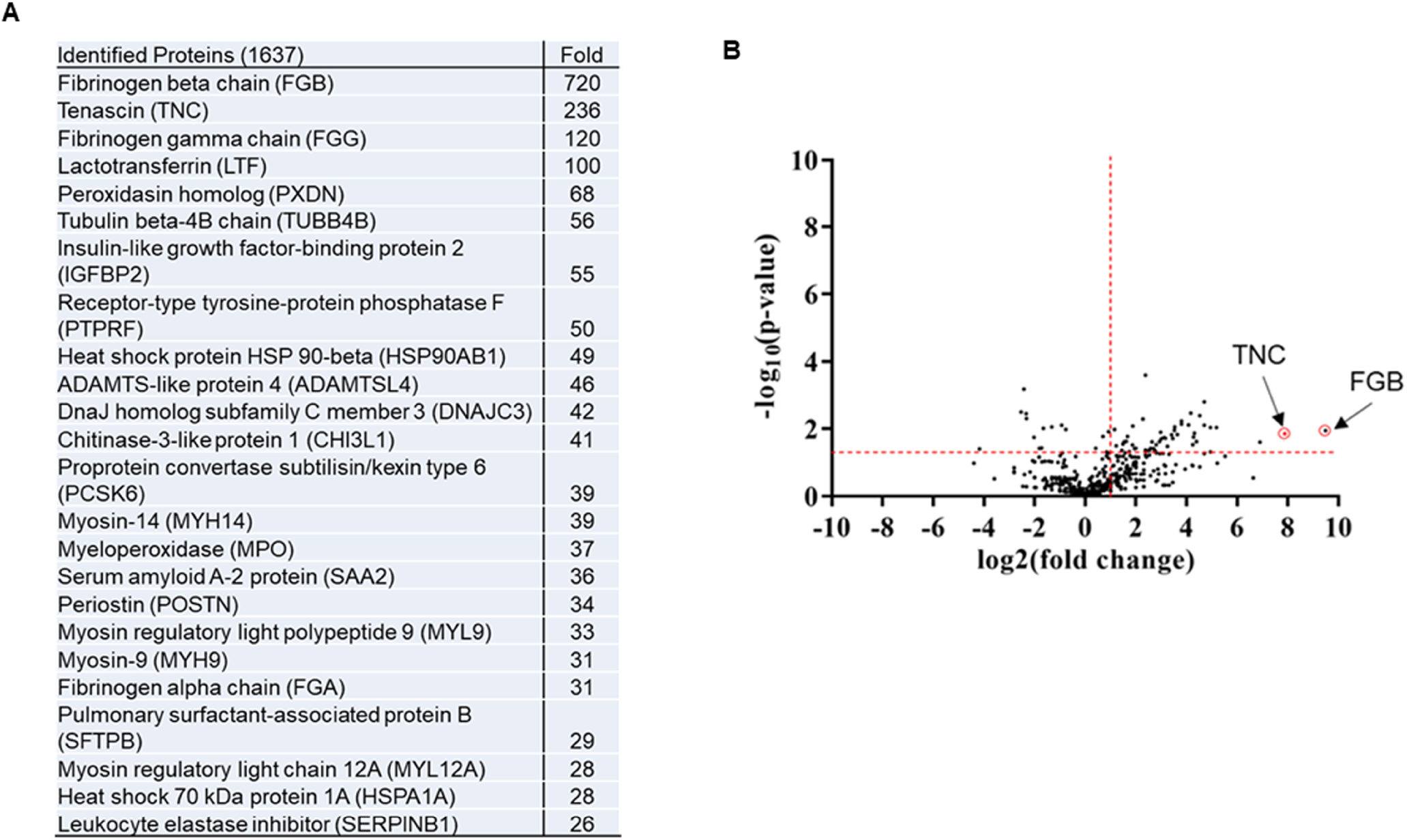
Comprehensive changes in plasma proteome profile of COVID-19 patients. **Panel A:** Top hits of COVID-19 exosomal proteins (fold) as compared to normal exosomes are shown as a fold increase. **Panel B:** Volcano plot illustrates significant difference in fold change of proteins in COVID-19 exosomes as compared to normal exosomes. The x-axis represents log2 - fold change and y axis is –log10 p-value showing statistical significance. Horizontal dashed red-line showing p =0.05 (–log10(0.05) = 1.3) and Vertical dashed red-line represents fold change (COVID/ normal exosomes) at 2 (log2(2) = 1). The absolute 2-fold change and p-value 0.05 were considered as the threshold cut-off. TNC and FGB are shown in red circles.

TNC is an immunomodulatory extracellular matrix glycoprotein that induces chronic inflammation and fibrosis in organs, including lung, liver, and kidney, by interaction with toll-like receptor 4 (TLR4) and integrin receptors (12). However, association of TNC in SARS-CoV-2 infection was not known. On the other hand, FGB is one of the components of fibrinogen complex cleaved by the protease thrombin into fibrin to form blood clot (13, 14). Increased levels of blood fibrinogen and associated disorders, such as coagulopathy and venous thromboembolism, are observed in COVID-19 patients (15). Enhanced expression of TNC and FGB in exosomes isolated from COVID-19 patients was verified by Western blot analysis **(Figure 3, panels A and B)**. However, the expression of these proteins in exosomes from normal plasma was significantly low. Acute inflammation is characterized by increased production of cytokines and the primary feature of COVID-19 is known to cause severe lung injury, and multi-organ failure (12, 16, 17). The FGB and TNC induce pro-inflammatory cytokine production through interaction with the inflammatory NF-kB signaling pathway and is supported by String analysis **(Figure 3, panel C)**. These results suggested COVID-19 patient plasma exosomes harbor TNC and FGB and transport to distant organs for virus associated pathogenesis.

**Figure 3:**
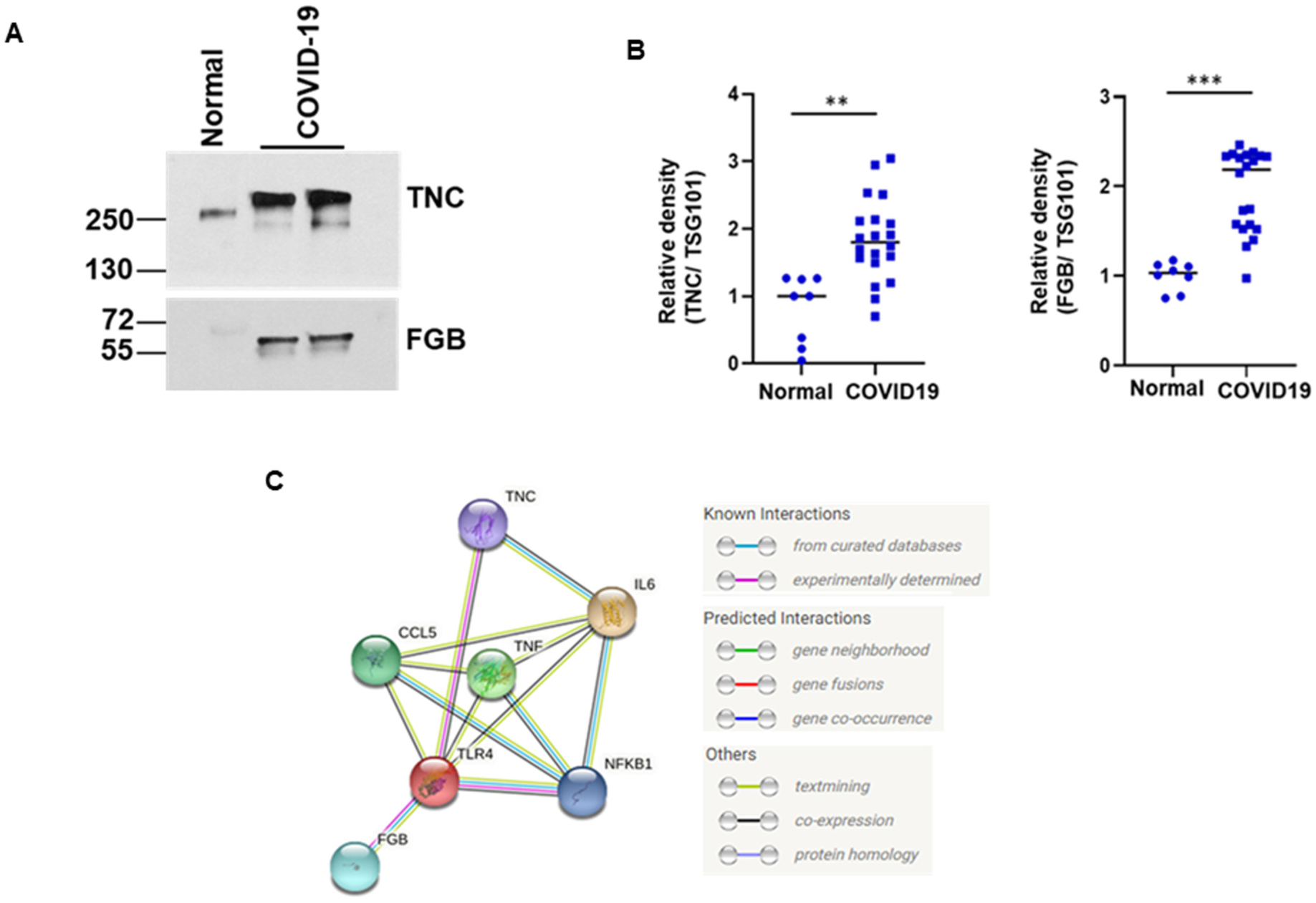
Tenascin-C (TNC) and fibrinogen-β (FGB) are highly present in exosomes of COVID-19 patients. **Panel A:** Lysates from COVID-19 plasma exosomes and control exosomes were subjected to Western blot analysis for TNC and FGB using specific antibodies and representative image is shown. **Panel B:** Dot plots for quantitative Western blot band intensities by densitometry analysis using ImageJ software (right panel) are shown (n=8 normal and n=20 COVID-19 samples). TSG101, an exosomal marker protein, was used for normalization of each sample. (**, p<0.01; ***, p<0.001). **C:** String analysis network module represents functional association of TNC and FGB with TLR4/NF-kB signaling. Each node represents all the proteins produced by a single protein coding gene locus. Colored node represents query proteins and first shell of interactions. Filled node shows 3D structure (known or predicted). Edges represent protein-protein associations for shared function.

### Exosomes isolated from COVID-19 plasma trigger pro-inflammatory cytokines in hepatocytes through activation of NF-kB signaling

To investigate the association, hepatocytes *in vitro* were used as a model cell line and exposed to exosomes from COVID-19 patients or from healthy controls. A significant upregulation of tumor necrosis factor-α (TNF-α), interleukin-6 (IL-6) and C-C motif chemokine ligand 5 (CCL5) were observed from exposure of immortalized human hepatocytes (IHH) to exosomes isolated from COVID19 patient plasma, as compared to exosomes from healthy normal plasma **(Figure 4, panel A)**. Similar results were noted from COVID-19 plasma exosomes when exposed to a different cell line with hepatocyte origin (Huh7). Pearson correlation analysis among expressions of the TNF-α, IL-6 and CCL5 in the hepatocytes exposed with patient exosomes. A significant positive correlation (P= 0.001, r= 0.66) was seen between TNF-α and CCL5 expression from hepatocytes **(Figure 4, panel B)**. These results demonstrated that COVID-19 plasma exosomes trigger strong pro-inflammatory cytokine production in hepatocytes.

**Figure 4:**
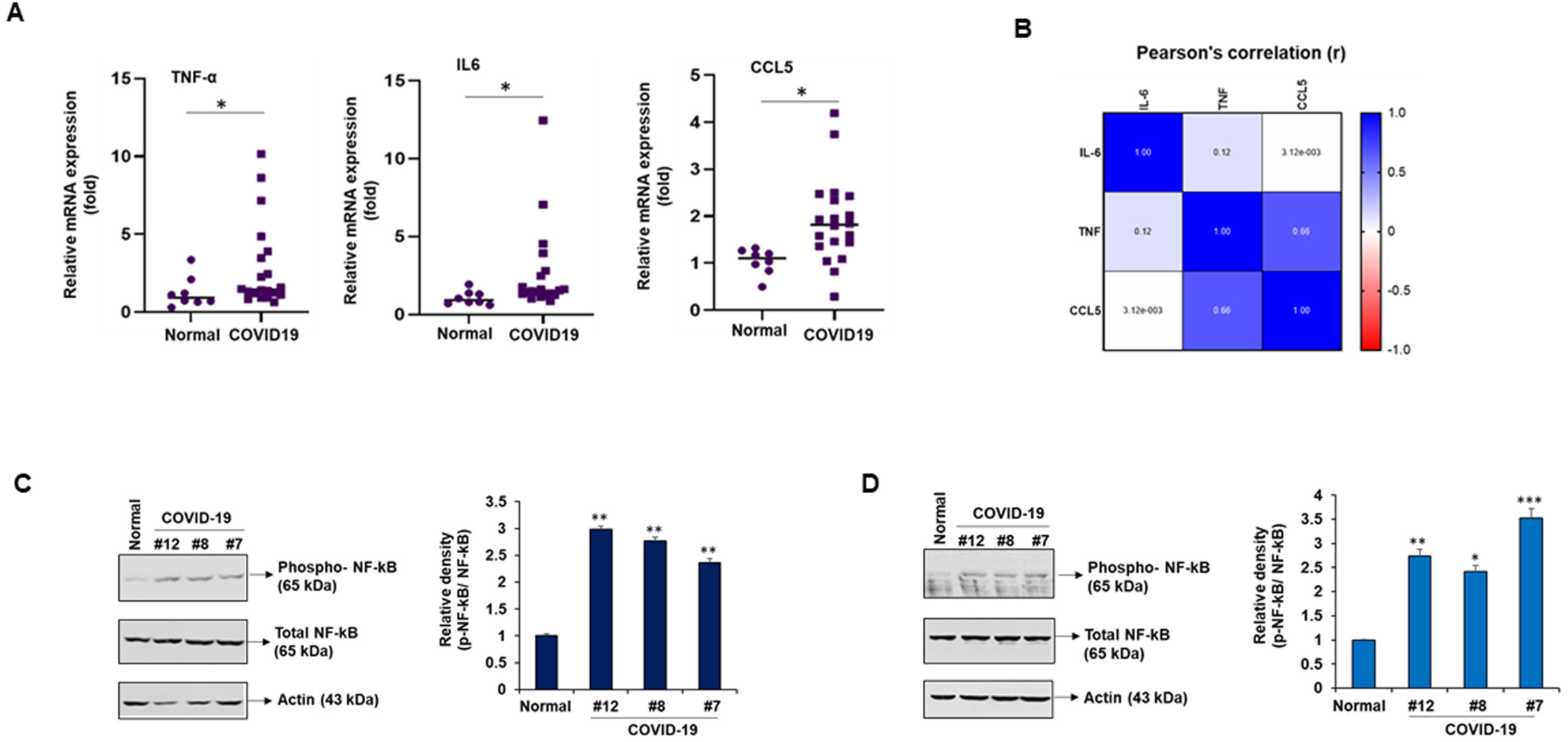
Exposure of hepatocytes to exosomes from COVID-19 plasma triggers pro-inflammatory molecules. **Panel A.** IHH were exposed to normal and COVID-19 exosomes for 48h and total RNA was isolated. Relative mRNA expression of TNF-α, IL-6 and CCL5 in cells treated with COVID-19 exosomes (n=20) or normal exosomes (n=8) were examined by qRT-PCR and represented by dot plots. 18s rRNA was used as an internal control. Small bar indicates standard error (* p<0.05). **Panel B:** Pearson correlation analysis among expressions of the TNF-α, IL-6 and CCL5 in the hepatocytes exposed with patient exosomes. **Panels C and D:** Exosomes from plasma of COVID-19 patients activates NF-kB signaling in hepatocytes. IHH (panel C) or Huh7 cells (panel D) were exposed with normal and COVID-19 exosomes for 48h and cell lysates were subjected to Western blot analysis for phospho-NF-kB p65 (Ser536), NF-kB p65 using specific antibodies. The membrane was reprobed for actin as an internal control. Right panel shows quantitative representation of Western blot band intensities. Small bar indicates standard error (*p<0.05; **p<0.01).

NF-κB is a key player that regulates inflammation, innate and adaptive immunity, proliferation, and cell survival (18). We observed a significant increase in phospho-NF-κB upon exposure of hepatocytes (IHH or Huh7) to exosome from COVID-19 patients as compared to that from normal plasma **(Figure 4, panels C and D)**. The results further demonstrated that exosomes from COVID-19 plasma, enriched with TNC and FGB, generate cytokine production by activation of NF-κB signaling in hepatocytes.

## Discussion

The novelty of observations from this study reveals: (i) an elevated level of TNC and FGB in exosomes from plasma of COVID-19 patients, and (ii) COVID-19 exosomes enhance expression of pro-inflammatory cytokines TNF-α, IL-6, and chemokine CCL5 through the NF-κB signaling pathway upon exposure to hepatocytes, as a model for cells from distant organ. Our results explain a different trans-regulatory mechanism for multiorgan pathogenic disorders during SARS-CoV-2 infection.

An array of clinical studies in the literature demonstrated that COVID-19 patients experience a cytokine storm (17, 19), although specific consequences and related mechanisms are not well defined. There are conflicting reports on the presence of SARS-CoV-2 RNA in patient blood, but we failed to detect viral RNA in the exosomes, and a similar observation was reported recently (3, 20). Hijacking the exosomal pathway by several RNA viruses has been demonstrated to mediate endogenous intercellular communication, immune modulation, and pathogenesis (6, 9, 21). We and others have shown that exosomes from HCV infected hepatocytes carry messages for activation of hepatic stellate cells and induces fibrosis marker expression (11, 22). GM3-enriched exosomes in COVID-19 patients are reported using lipidomics (23), although the functional consequence is yet to be determined.

We focused at this point on the inflammatory molecules since they play a major role in COVID-19 related pathogenesis. The association of TNC with SARS-CoV-2 infection has not been previously reported. TNC induces chronic inflammation and fibrosis via an interaction with toll-like receptor 4 and integrin receptors. Elevated expression of TNC in exosomes is reported in glioblastoma patients and in nasal lavage fluid during human rhinoviruses infection (12, 24). On the other hand, increased level of blood fibrinogen and venous thromboembolism are reported in COVID-19 patients (15). In addition, FGB is involved in other disease processes, such as wound healing, liver injury, allergic airways disease, cardiovascular disease and microbial pathogenesis by modulating host immune system. FGB is primarily synthesized in hepatocytes, although extrahepatic epithelial cells also synthesize fibrinogen (25). Elevated FGB expression was reported in lung adenocarcinoma and suggested a potential role as biomarker (26). Elevated fibrinogen is detected in exosomes from drug and alcohol induced liver injury, neurological disorder, and swine flu viral infection (14, 27–30).

Chronic inflammation and increased cytokine production, like TNF-α, IL-6 and CCL5, are critical features of COVID-19 patients which may suggest causing severe lung injury, multi-organ failure, and poor prognosis (16, 17, 31–33). We observed activation of NF-KB signaling and activation of target genes, TNF-α, IL-6 and CCL5, following exposure of hepatocytes to exosomes isolated from plasma of COVID-19 patients **(Figure 5)**. In SARS-Co V2 infection, increased level of IL-6 and STAT3 are reported (31, 34). The spike protein of SARS-Co V2 triggers IL-6 production (5) and analyzing SARS-CoV-2 genetic materials in exosomes may be important in understanding whether viral genetic materials play a role in cytokine induction in distant organs. Other molecules present in exosomes from plasma of COVID-19 patients may also have role in disease processes and need to be evaluated in future studies. Several studies have suggested that non-coding RNAs carried through the exosomes play a crucial role in virus mediated disease progression and warrant investigation. The presence of TNC and FGB in SARS-CoV-2 infected patient exosomes may be a mechanism for cytokine storm resulting in micro-thrombosis in some patients. Exosomes used in this study were isolated from patients admitted to ICU in our academic medical center. Analyzing exosomes from SARS-CoV-2 infected patients with mild symptoms in future will help in understanding whether TNC and/or FGB can be used as the potential prognostic markers. Further, how TNC and FGB are enhanced COVID-19 patient will be important to understand. In conclusion, our results suggested that exosomes carry TNC and FGB in hospitalized COVID-19 patients, and exposure of cells from distant organs may trigger cytokine expression. Our work also highlighted that TNC and FGB-enriched exosomes from COVID-19 plasma and may be correlated for the first time with pathogenesis.

**Figure 5:**
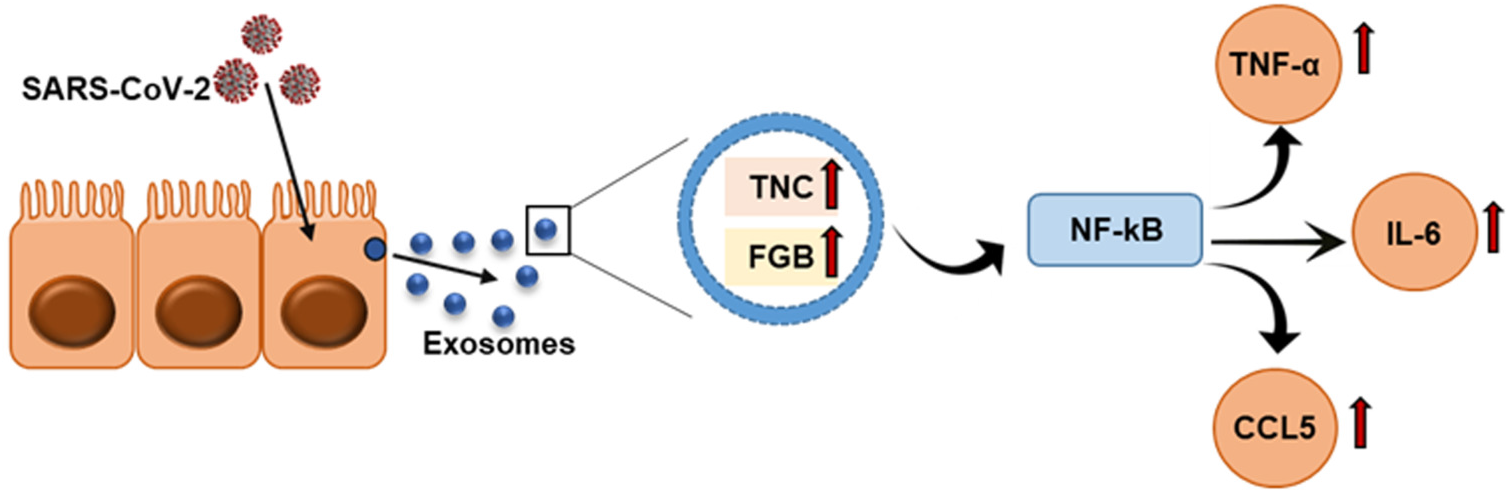
Schematic representation shows exosomes secreted from SARS-CoV-2 infected cells are enriched with TNC and FGB triggering TNF-α, IL-6 and CCL5 production in hepatocytes via NF-κB signaling.

## Materials and Methods

### Plasma specimens

A total of 20 deidentified, heparinized-plasma specimens collected from COVID-19 patients admitted at the Saint Louis University Hospital, were used. All patients were confirmed positive for SARS-CoV-2 by RT-PCR performed on a nasopharyngeal swab around the time of hospital admission. Patient specimens were collected for clinical laboratory analyses as part of routine clinical care. Patient information is summarized in Table 1. This study was waived by the Saint Louis University Institutional Review Board for use of de-identified clinical specimens. Archived plasma samples from 8 healthy volunteers were included as normal control and were collected from pre-COVID-19 era for a different study.

### Exosome isolation and analysis

Exosomes were isolated from plasma using the ME Kit (ME-020p-Kit) following the supplier’s instruction (New England Peptide Inc, MA). The exosomes were examined after negative staining using a JEOL JEM-1400Plus Transmission Electron Microscope.

### Cell culture and exposure with exosomes

Immortalized human hepatocytes (IHH) and a human hepatoma cell line (Huh7) were maintained in Dulbecco’s modified Eagle’s medium (DMEM) supplemented with 10% fetal bovine serum (FBS) and 1% penicillin/streptomycin at 37°C in a 5% CO_2_ atmosphere. IHH and Huh7 cells were seeded into 6-well plate at a density of 3×10^5^ cells/well and exposed to equal concentration of exosomes for 48h. Cells were harvested for RNA analyses.

### Mass spectrometry analysis

The exosome pellet was dissolved in lysis buffer (4% SDS, 100 mM DTT, 150 mM Tris–HCl pH 8.0) was subjected to Mass Spectrometric analysis using a Thermo Q-Exactive system. Peptides were separated on an EASYnLC system with a Thermo ES803 PepMap C18 column. The results were acquired in data dependent acquisition mode; top10 m/z for MS2 per cycle (Washington University Proteomics Shared Resource). Candidate proteins were defined as those have minimum 5 spectrum count and at least 2-fold enrichment compared to normal.

### RNA isolation and analysis

Total RNA was isolated from exosomes or hepatocytes (IHH or Huh7) for qRT-PCR as described previously (11), using TaqMan Universal PCR master mix and 6-carboxyfluorescein (FAM)-MGB probes for SARS-CoV-2 [2019-nCoV CDC EUA kit (10006770, IDT)], CCL5(assay ID: HS009822282_m1), IL-6 (assay ID: HS00985639_m1) and TNF-α: (assay ID: HS00174128_m1) following manufacture’s protocol (Thermo Fisher Scientific). 18s (assay ID: Hs03928985_g1) was used as endogenous control. The relative gene expression was analyzed by using the 2^-ΔΔCT^ formula (*ΔΔC_T_* = *ΔC_T_* of the sample – *ΔC_T_* of the control). Each sample was loaded in triplicate for analysis.

### Western blot analysis

Cells lysates were subjected to Western blot analysis using specific antibodies to CD63 (Santa Cruz Biotechnology), TSG101 (Santa Cruz Biotechnology), tenascin (TNC) (Sigma), fibrinogen-β (FGB) (Santa Cruz Biotechnology), phospho-NF-kB p65 (Ser536) (Cell Signaling Technology, CST), NF-kB p65 (CST). The blot was reprobed with actin-HRP antibody (Santa Cruz Biotechnology) to compare protein load in each lane. Densitometry analysis was done using Image J software.

### Statistical Analysis

The results were expressed as mean ± standard error. Student′s t test was used for comparison between two groups (normal vs. COVID-19 exosomes). Pearson’s correlation analysis was performed using GraphPad Prism software. P-values of <0.05 were considered statistically significant. All experiments were repeated at least three times, and representative data are shown.

## Acknowledgements

Our research was supported by the Research Institute of Saint Louis University (T.S.I and R.R) and Pathology Department Seed Grant (RBR). The funders had no role in study design, data collection and analysis, preparation of the manuscript, or decision to publish.

## Conflict of Interest

No potential conflict of interest is disclosed.

